# Togaram Ensures Axial Alignment of the Sperm Neck

**DOI:** 10.64898/2026.04.15.718719

**Authors:** Emma E. Burns, Sara E. Burr, Kathleen H. M. Holmes, Carey J. Fagerstrom, Nasser M. Rusan

## Abstract

The sperm neck must remain mechanically stable to preserve a straight head-tail axis during spermiogenesis, yet the mechanisms that maintain this alignment after head-tail attachment forms are poorly understood. Here, using *Drosophila*, we identify Togaram (Toga), the ortholog of mammalian TOGARAM1, as a regulator of this maintenance phase. Toga localizes to the sperm neck and to a broader perinuclear microtubule scaffold that surrounds the nucleus, which we term the Sperm Microtubule Cage (SMC). Depletion of Toga causes a striking off-axis phenotype in which the centriole usually remains attached to the nucleus but fails to maintain a straight head-tail axis, whereas complete detachment is comparatively rare. Live imaging shows that this defect arises after initial attachment has formed, indicating a maintenance defect rather than a failure of head-tail assembly. Mechanistically, Toga is dispensable for initial formation of head and neck microtubules but is required for their stabilization during Canoe-stage remodeling. Loss of Toga increases microtubule turnover, strongly reduces acetylated tubulin in the manchette, SMC, and neck region, and leaves the sperm neck vulnerable to buckling. Off-axis spermatids are depleted at later stages, consistent with selective loss of mechanically compromised cells. Finally, Cep104 and CCDC66 associate with Toga in cultured cells and are likewise required for axial alignment, consistent with a conserved module analogous to the mammalian ciliary tip module. Together, these findings support a model in which Toga stabilizes sperm head and neck microtubules to preserve a straight head-tail axis during the maintenance phase of spermiogenesis.

## INTRODUCTION

The transformation of a round spermatid into highly polarized, motile sperm is a dramatic example of cellular remodeling. During spermiogenesis, a post-meiotic cell coordinates nuclear condensation and reshaping, axoneme elongation, and formation of a robust connection between the sperm head and tail. This connection is mediated by the Head-Tail Coupling Apparatus (HTCA), a specialized composite of nuclear-envelope, centriolar, and cytoplasmic proteins and accessory structures (Buglak et al., 2025). Across diverse species, including humans, mice, bovine, and flies, disruption of HTCA assembly or maintenance causes male infertility through defective head-tail coupling, often culminating in partial or complete separation of the head from the tail (Blom and Birch-Andersen, 1970; Buglak et al., 2024; Gan et al., 2024; Hetherington et al., 2016; Kracklauer et al., 2010; Shang et al., 2018; Shang et al., 2017; Shen et al., 2020; Texada et al., 2011; Texada et al., 2008; Wu et al., 2024; Yang et al., 2018; Yang et al., 2012; Yuan et al., 2015; Zhang et al., 2021a; Zhang et al., 2024; Zhang et al., 2021b). In humans, these phenotypes are broadly represented within Acephalic Spermatozoa Syndrome, which not only include complete decapitation but can also include severe head-tail misalignment, emphasizing that physical attachment and axial alignment are separable but equally essential features of a functional sperm (Baccetti et al., 1989; Chemes et al., 1999; Chemes and Rawe, 2010; Perotti et al., 1981).

In *Drosophila*, the initial establishment of the head-tail linkage occurs in early spermatids shortly after meiosis II and is dependent on microtubules and dynein-mediated transport of the centriole to the nuclear envelope (Galletta et al., 2020; Li et al., 2004). Once established, this linkage must be maintained throughout the extensive structural remodeling of later spermiogenesis. This maintenance phase presents a significant mechanical challenge, as the connection must withstand the forces generated by axoneme elongation and dramatic nuclear reshaping (Fabian and Brill, 2012; Tokuyasu, 1975). Furthermore, it is important to explicitly note that attachment and alignment are not synonymous, as a centriole and nucleus can remain connected yet still fail mechanically by buckling or bending under developmental strain (Galletta et al., 2020). To achieve this stability, several mechanisms must function in coordination.

First, the HTCA provides the adhesive framework that anchors the centriole to the nuclear envelope, utilizing components such as the SUN-domain protein Spag4 and other specialized factors (Buglak et al., 2024; Kracklauer et al., 2010; Texada et al., 2008). Second, accessory structures such as the striated columns and capitulum likely provide architectural reinforcement (Buglak et al., 2025). Finally, the sperm neck, defined as the entire region between the axoneme and the nucleus (which includes the centriole), must remain sufficiently rigid to preserve a linear head-tail axis while accommodating tissue remodeling. Despite the importance of this region, the molecular mechanisms that confer neck stability and prevent buckling under developmental strain remain poorly understood.

Microtubules are strong candidates to provide this missing mechanical resilience. Around the sperm head, perinuclear microtubule arrays contribute to nuclear shaping and likely serve as substrates for force transmission during elongation (Fawcett et al., 1971; Li et al., 2023; Riparbelli et al., 2020). The best-known example is the manchette, a transient microtubule-dense scaffold that surrounds the nucleus and helps drive sperm head remodeling (Fawcett et al., 1971; Li et al., 2023; Riparbelli et al., 2020). In Drosophila, Mst27D physically links nuclear pore complexes to bundled microtubules during nuclear elongation, underscoring the idea that head-associated microtubules are active structural elements rather than passive bystanders (Li et al., 2023). These observations suggest that successful spermiogenesis depends not only on building perinuclear microtubule arrays, but also on stabilizing them at the right place and time. However, how these microtubules are reinforced at the sperm neck to preserve axial alignment remains unknown.

In this study, we identify Togaram (Toga) as a critical regulator of sperm axial alignment. We show that Toga localizes to the sperm neck and a broader circum-nuclear scaffold we term the Sperm Microtubule Cage (SMC), which encompasses the manchette. Our data indicate that Toga is required after initial head-tail attachment has formed, where it promotes microtubule stabilization and acetylation to prevent buckling of the sperm neck during Canoe-stage remodeling. We further show that Cep104 and CCDC66 share this axial-alignment function and can associate with Toga, consistent with a conserved module analogous to the mammalian Ciliary Tip Module (CTM). Together, these findings support a model in which Toga stabilizes sperm head/neck microtubules to preserve a straight head-tail axis during the maintenance phase of spermiogenesis.

## RESULTS AND DISCUSSION

### Togaram is required to maintain sperm axial alignment

While the critical nature of the Head-Tail Coupling Apparatus (HTCA) in spermatogenesis is well established, the molecular mechanisms that preserve sperm neck alignment during the maintenance phase of spermiogenesis remain incompletely defined. To identify unknown proteins that localize to the sperm neck, we screened GFP trap lines of candidate proteins. We selected proteins identified in the *Drosophila* sperm proteome (Garlovsky et al., 2022; Wasbrough et al., 2010) and prioritized both uncharacterized and cytoskeleton binding proteins. Our initial screen of 30 proteins relied on a simple qualitative analysis of protein localization in spermatids (Table S1).

Our analysis identified one protein, CG42399, that localized at the sperm neck (Fig. 1A). CG42399 is the ortholog of the mammalian TOGARAM1, showing 38% amino acid similarity and 22% identity (Ozturk-Colak et al., 2024). TOGARAM1 is a Tumor Overexpressed Gene (TOG)-domain containing protein within the Crescerin family, which has been shown to regulate axonemal microtubules (Das et al., 2015; Latour et al., 2020). In *C. elegans*, the ortholog CHE-12 ensures sensory cilia integrity, suggesting a deep evolutionary conservation of this family in stabilizing microtubule-based organelles (Bacaj et al., 2008; Das et al., 2015). Based on this homology, we refer to the CG42399 *Drosophila* gene as *togaram* (*toga*).

**Figure 1:**
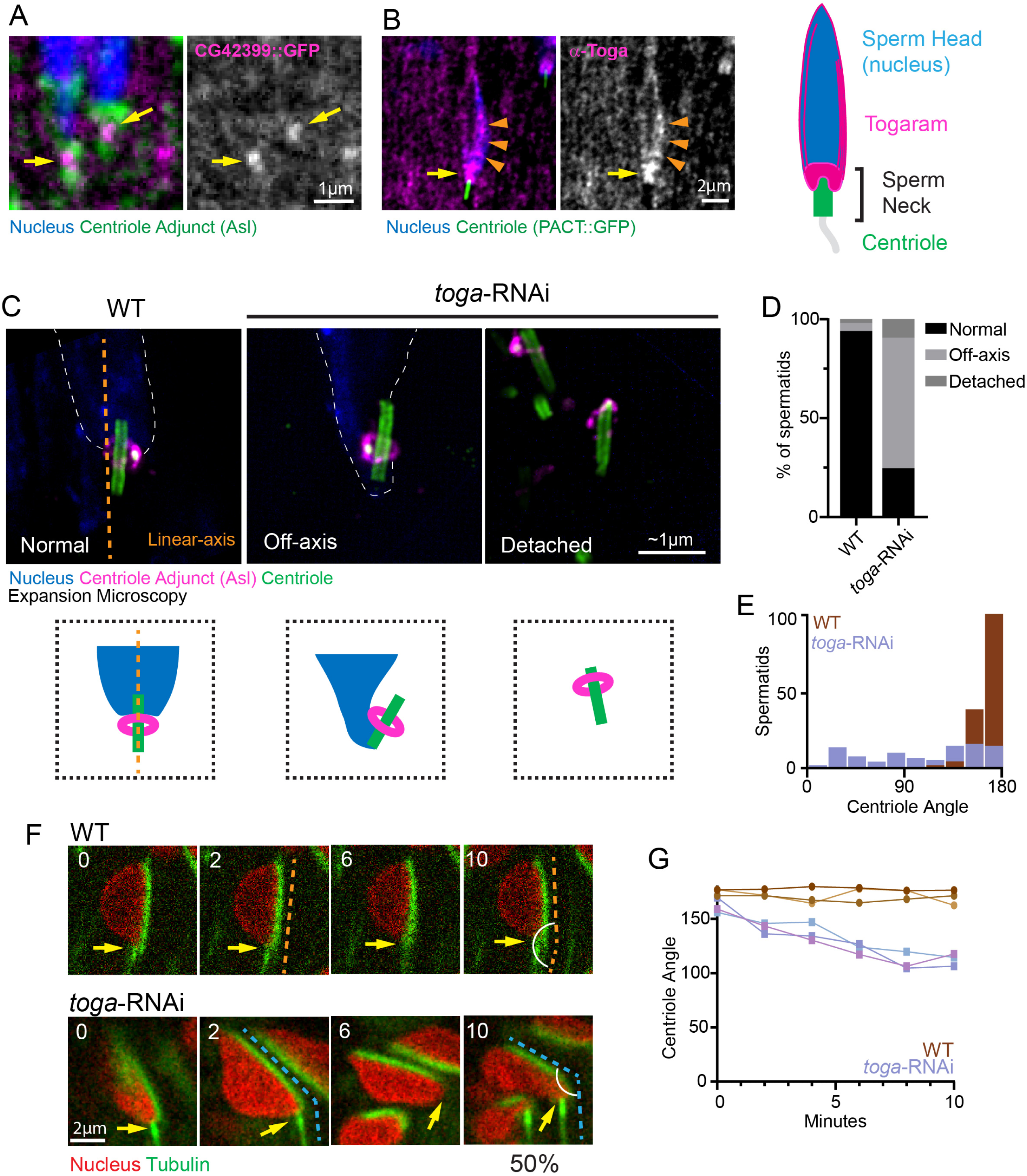
Togaram is required for sperm axial alignment. (A) Representative image showing Canoe stage spermatid showing Togaram (CG42399::GFP, magenta), the Centriole Adjunct (CA; Asterless staining, green) and the nucleus (DAPI, blue). Togaram neck localization (yellow arrows). Scale bar: 1 μm. B) Representative image of Togaram immunofluorescence (Toga, magenta), the centriole (PACT::GFP, green), and the nucleus (DAPI, blue) in a Canoe stage spermatid. Toga neck localization (yellow arrows); Toga head localization (orange arrowheads). Cartoon of Canoe stage spermatid showing Toga (magenta) neck and head localization. Scale bar: 2 μm. (C) Representative Expansion Microscopy (ExM) images showing wild-type (left) and *toga*-RNAi (right two images) Canoe stage spermatids. Spermatids were labeled for the nucleus (DAPI, blue, dotted line), the CA (Asl, magenta), and the centrioles (Ana1, green). Cartoons depict spermatids with centrioles that are normal, off-axis, or detached in relation to nucleus. Dotted orange line shows linear alignment of normal spermatids. Scale bar: ∼1 μm. (D) Quantification of wild-type (n=150) and *toga*-RNAi (n=76) spermatids alignment phenotypes. (E) Quantification of attached centriole angle relative to nucleus in wild-type (n=150) and *toga*-RNAi (n=76). (F) Representative frames of live imaging wild-type (n=6) and *toga*-RNAi (n=6) cysts of spermatids labeled for the nucleus (H2A::RFP) and microtubules (ubi-GFP::tubulin). Yellow arrows indicate spermatid neck region. Dashed lines denote proper alignment (orange) and off-axis misalignment (blue). Scale bar: 2 μm. (G) Quantification of changes in the centriole angle over time from representative wild-type and *toga*-RNAi movies.

To confirm localization to the neck, we raised an antibody against a Toga fragment spanning amino acids 1–664. Consistent with the endogenously tagged (intronic insertion gene trap) GFP::Toga (Nagarkar-Jaiswal et al., 2015), immunofluorescence staining confirmed Toga localization to the sperm neck region (Figure 1B, Figure S1, yellow arrows). However, the antibody revealed an additional localization of Toga along the elongating nucleus from Early to Late Canoe stage spermatids (Figure 1B, Figure S1, orange arrowheads). This second localization is intriguing given that nuclear reshaping in *Drosophila* largely relies on a perinuclear, microtubule-dense manchette localized along the nucleus (Fawcett et al., 1971; Li et al., 2023; Riparbelli et al., 2020). These distinct localization pools suggest Toga performs multiple functions in spermatids, potentially coordinating nuclear remodeling with HTCA assembly.

Given the localization of Toga to the sperm neck, we hypothesized that Toga plays a role in establishing and/or maintaining the head-tail connection in spermatids. To test this, we depleted Toga in the male germline using *bam-Gal4* to drive *uas-toga*-RNAi (Dietzl et al., 2007). In wild-type Early Canoe stage spermatids, the centriole attaches to the nucleus end-on, essentially forming a single linear axis from the axoneme, through the neck region and into the nucleus (Figure 1C,D; orange dotted line). By contrast, we found that approximately 65.8% of centrioles in *toga-*RNAi spermatids were misaligned (off-axis); an additional 9.5% of centrioles were completely detached from nuclei (Figure 1C,D). To better characterize the off-axis phenotype, we assessed the angle of the centriole relative to the nucleus. Wild-type spermatids had a consistent centriole attachment angle of approximately 180 degrees with little variation (Figure 1E). However, following *toga-*RNAi, the angle of centriole attachment to the nucleus was highly variable with nearly 50% forming an acute angle (Figure 1E). These data suggest Toga plays a pivotal role in the stability of the neck during spermiogenesis.

This “off-axis” phenotype differs from the complete head detachment, or decapitation, observed in mutants such as *Spag4* (a SUN-domain protein) (Kracklauer et al., 2010). Instead, the *toga* phenotype is primarily a misalignment phenotype, suggesting that the fundamental adhesive connection between the centriole and nucleus, the HTCA itself, remains intact but lacks structural rigidity. In fact, we show that *toga-*RNAi does not have an effect on Spag4 localization (Figure S2, yellow arrows).

Since mammalian TOGARAM1 promotes microtubule polymerization and bundle stability (Das et al., 2015; Latour et al., 2020; Saunders et al., 2025), we propose that Toga stiffens the microtubule interface between the basal body and the nucleus, otherwise stabilizing the neck. We hypothesize that without Toga, the sperm neck becomes flexible and susceptible to buckling, unable to resist the surrounding mechanical forces of the entire tissue, thus failing to maintain the spermatid axial alignment.

While our analysis of fixed tissue revealed an alignment defect, it remained unclear whether the phenotype arose from a failure in initial attachment or a progressive loss of stability. To investigate the temporal dynamics of the off-axis defect, we performed live imaging of explanted testes expressing *H2A::RFP* to mark nuclei and *ubi-GFP::tubulin* to visualize the microtubules. In wild-type cysts, we observed a stable alignment between the sperm head and the growing axoneme (Figure 1F, G). In contrast, *toga*-RNAi spermatids frequently exhibited misalignment. In many cases, we captured properly aligned spermatids that, over time, appeared to bend at the HTCA (Figure 1F, G). These movies support our mechanical stability model whereby the head-tail attachment forms correctly in spermatids lacking Toga, but yields under the mechanical strain of tissue remodeling during development. Thus, Toga appears to be required for the maintenance, rather than the initial establishment, of the sperm neck axis.

### Spermatids with off-axis centrioles are likely eliminated during spermiogenesis

Interestingly, fixed analysis revealed that later stages of spermiogenesis, the Mid-and Late Canoe stages, exhibited proper axial alignment of the sperm head and neck in the majority of spermatids (Figure 2A, orange dotted line). This apparent recovery was unexpected given the severity of the defects observed in earlier stages (Figure 2A, blue dotted line; Figure 1C,D). We proposed two hypotheses: either the off-axis centrioles eventually correct their alignment, potentially aided by an intact, albeit flexible, HTCA, or alternatively, *toga*-deficient spermatids that fail to maintain alignment are segregated and ultimately eliminated.

**Figure 2:**
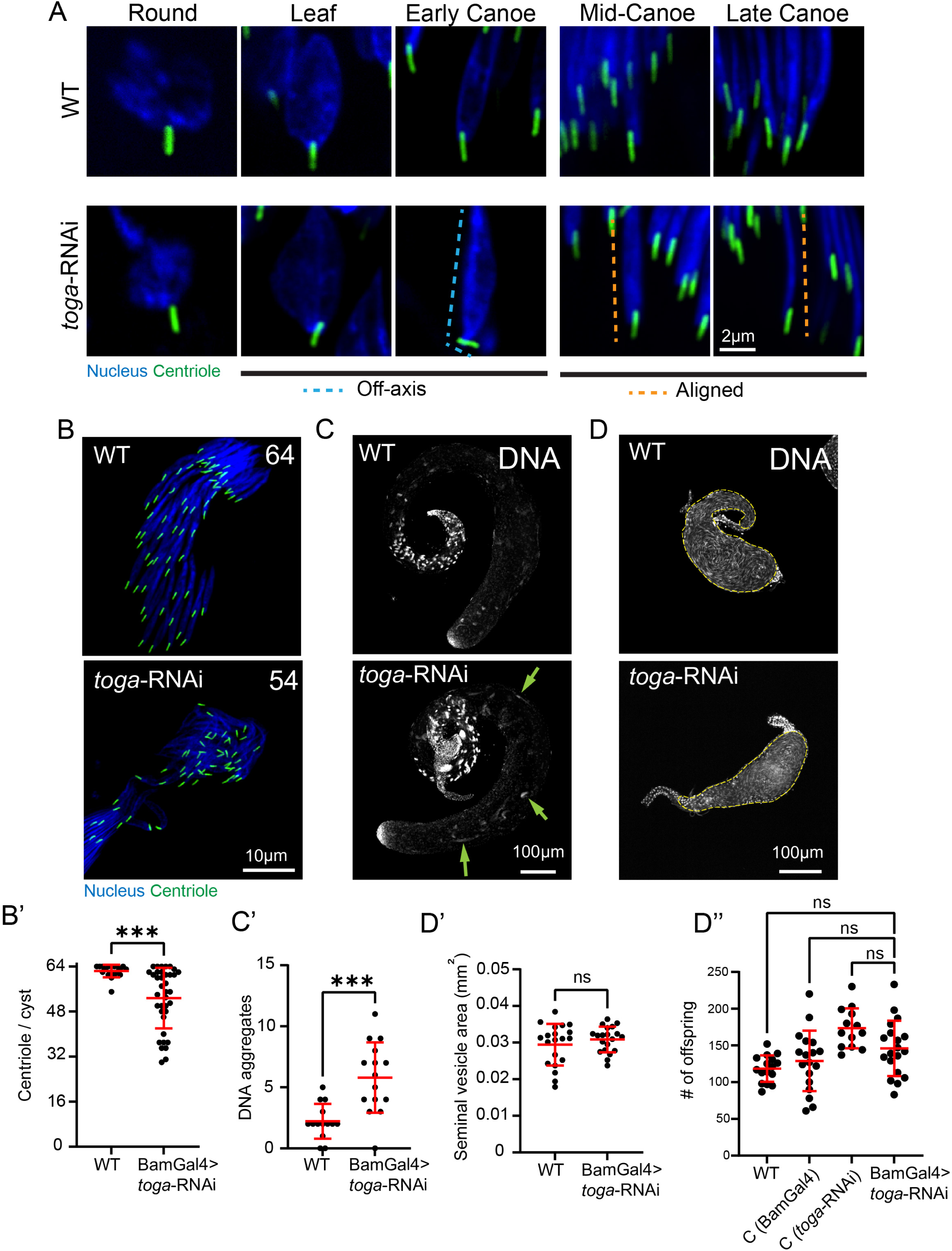
Off-axis spermatids are likely eliminated late in spermiogenesis. (A) Representative images showing wild-type (top) and *toga*-RNAi (bottom) spermatids during indicated developmental stages. Spermatids are labeled for the nucleus (DAPI, blue) and the centriole (PACT::GFP, green). Scale bar: 2 μm. (B) Representative images showing wild-type (top) and *toga*-RNAi (bottom) cysts of spermatids labeled for the nucleus (DAPI, blue) and the centriole (PACT::GFP, green). Number indicates number of centrioles in representative cysts. Scale bar: 10 μm. (B’) Quantification of wild-type (n=19) and *toga*-RNAi (n=37) centrioles per cyst. (C) Representative images showing wild-type (top) and *toga*-RNAi (bottom) testes labeled for the DNA (DAPI). Green arrows indicate DNA aggregates. Scale bar: 100 μm. (C’) Quantification of wild-type (n=14) and *toga*-RNAi (n=14) mislocalized DNA aggregates per testis. (D) Representative images showing wild-type and *toga*-RNAi seminal vesicles (dashed yellow line). Scale bar: 100 μm. (D’) Quantification of seminal vesicle area in wild-type (n=20) and *toga*-RNAi (n=20) testes. (D’’) Quantification of offspring from control group males: yw (n=16), BamGal4/+ (n=18), *toga*-RNAi/+ (n=13) and experimental group males: BamGal4>*toga*-RNAi (n=19). Error bars are mean±s.d., ns = not significant, \*\*\**P*≤0.0003 (unpaired two-tailed t-test for B’, C’, D’, one-way ANOVA with Tukey’s correction for E).

To distinguish between axis-correction and spermatid elimination, we counted the number of centrioles in wild-type and *toga*-RNAi cysts. As spermatids exit meiosis, the 64 nuclei and 64 nuclei-associated centrioles are present as a tight cluster at one end of each cyst. In. In contrast to wild-type cysts that contain an average of 62 centrioles, *toga* RNAi cysts contained an average of only 53 (Figure 2B,B’). While we were unable to locate and precisely count all the centrioles and nuclei along the entire length of cysts, we did detect many misplaced nuclei and abnormal DNA aggregates, and some centrioles, throughout toga RNAi testes (Figure 2C, C’). These observations are consistent with elimination of defective sperm. This reduction supports a model whereby the loss of Toga compromises sperm neck integrity to an extent that prevents development beyond the Early Canoe stage. Spermatids that suffer severe mechanical failure are likely depleted, leaving only the mechanically sound spermatids to progress.

The observed “fall out” phenotype is reminiscent of defects in sperm bundle organization involving the Head Cyst Cell (HCC), which normally anchors maturing spermatid heads in a tight cluster (Desai et al., 2009; Lindsley, 1980; Tokuyasu et al., 1972). One possible model is that severe neck buckling prevents some toga-deficient spermatids from remaining stably engaged with the HCC niche. This idea remains speculative, as we do not directly visualize HCC disengagement here, but it is consistent with the reduced centriole counts, ectopic DNA aggregates, and the precedent that spermatids with defective nucleus-cytoskeleton coupling can become scattered or lost from the bundle, as in *yuri* mutants (Texada et al., 2008). Thus, the apparent recovery of alignment in late stages is likely a result of the system filtering defective sperm, ensuring that only structurally intact cells persist to the sperm individualization stage.

Based on the high number of spermatids that do survive the elimination stage in *toga*-RNAi, we predicted male fertility to be unaffected. Indeed, when we inspected seminal vesicle area as a proxy for sperm number, and performed a fertility assay, we detected no significant difference from controls (Figure 2D, D’, D’’). One might conclude that Toga is therefore not essential for lineage maintenance; however, it is possible that small changes in fecundity not detected in a lab assay might still exert profound negative effects in competitive, natural environments where subtle structural defects can significantly impact reproductive fitness.

### Togaram stabilizes sperm head and neck microtubules

To understand the mechanistic basis of head-tail axial-alignment defect, we considered whether Toga regulates sperm head and neck microtubules. This hypothesis was motivated by the presence of three TOG domains in Toga, a motif known to bind microtubules (Slep, 2009), and by prior work showing that mammalian TOGARAM1 stabilizes ciliary microtubules (Das et al., 2015; Latour et al., 2020; Saunders et al., 2025). TOGARAM1 appears to confer microtubule stability through three nonexclusive mechanisms. First, TOGARAM1 caps microtubule plus-ends by forming a cork-like structure as a member of the Ciliary Tip Module (CTM), which acts as a “slow polymerase” that promotes processive elongation and limits catastrophe (Saunders et al., 2025). Second, TOGARAM1 can bind the lattice of microtubules, which drives microtubule bundling as shown in overexpression experiments (Das et al., 2015). Third, TOGARAM1 promotes microtubule acetylation and polyglutamylation. This is likely indirect as TOGARAM1 is not a tubulin modifying enzyme, but is thought to promote the stable microtubule substrate required for the accumulation of these modifications (Latour et al., 2020; Saunders et al., 2025). Thus loss of Toga results in overall loss of microtubule stability and post-translational modifications, which has downstream consequences on cilia form and function (Bosch Grau et al., 2013; Shida et al., 2010; Wloga et al., 2009; Yang et al., 2014).

To test the hypothesis that Toga regulates microtubules that surround the nucleus and sperm neck, we first examined microtubule density using quantitative fluorescence analysis to measure GFP::tubulin levels as a proxy for the number of microtubules. We used live imaging for this analysis to avoid variation in fixation, extraction, and staining against tubulin. Imaging proved to be extremely difficult given the tissue density, but we were able to compare the average microtubule fluorescence surrounding sperm heads. While the average fluorescence intensity was reduced in the *toga*-RNAi condition (Figure 3A, A’), the data suggest that the off-axis phenotype is not caused by a catastrophic loss of microtubules. While microtubule polymer level was not drastically changed, we hypothesized that there might be an underlying change in microtubule dynamics or stability surrounding the head and neck. To test this hypothesis, we performed fluorescence recovery after photobleaching (FRAP) of spermatids expressing GFP::tubulin under the ubiquitin promoter (*ubi-*). We then measured the extent of fluorescence recovery to infer the relative fraction of dynamic versus stable microtubules. In Early Canoe stage spermatids, we found that following FRAP, both wild-type and *toga* RNAi spermatids recovered nearly identical amounts of tubulin during the course of the experiment (28% and 30%, respectively; Figure 3B-B’’). This suggests that the initial assembly of perinuclear microtubules in early spermiogenesis proceeds independently of Toga. However, as spermiogenesis progresses, the manchette undergoes significant stabilization through enhanced crosslinking to the nucleus, and likely other mechanisms (Fabian and Brill, 2012). Consistent with this, we observed a decrease in microtubule turnover in Mid-canoe stage wild-type spermatids, with less tubulin recovering than in Early Canoe, (mobile fraction dropping to 11% vs 28%, respectively). This shift marks a critical transition from a dynamic, remodeling cytoskeleton to a rigid, structural scaffold. This stabilization was slightly compromised in *toga*-RNAi spermatids where the average microtubule recovery was higher at 20% (Figure 3C-C’’). This indicates that, without Toga, microtubules are more dynamic, potentially failing to achieve their mature stable state.

**Figure 3:**
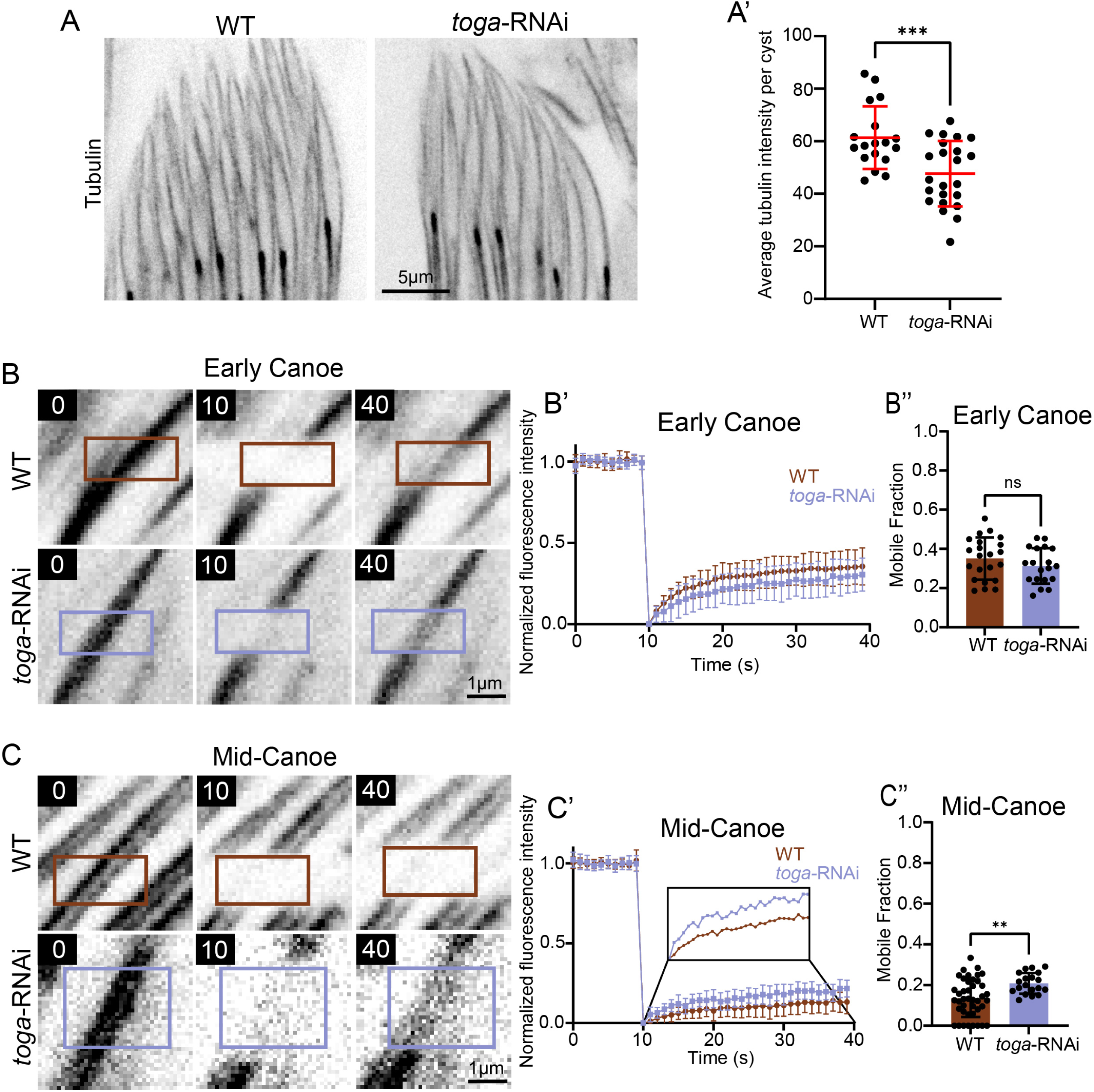
Togaram stabilizes microtubules at the sperm head. (A) Representative images showing wild-type (left) and *toga*-RNAi (right) spermatids labeled for tubulin (ubi-GFP::tubulin). Scale bar: 5 μm. (A’) Quantification of wild-type (n=18) and *toga*-RNAi (n=23) average tubulin intensity of sperm heads per cyst. (B) Representative frames of Early Canoe stage wild-type (top) and toga-RNAi (bottom) FRAP experiments of tubulin at sperm head before bleaching, immediately after bleaching (10 sec), and 30s after of recovery (40 sec). Boxes indicate photobleached area. Scale bar: 1 μm. (B’) FRAP graphs of Early Canoe stage wild-type (n=23) and *toga*-RNAi (n=19) spermatid. (B’’) Quantification of data in B’. (C) Representative frames of Mid-Canoe stage wild-type (top) and *toga*-RNAi (bottom) FRAP experiments of tubulin at sperm head before bleaching, immediately after bleaching (10 sec), and 30s after of recovery (40 sec). Scale bar: 1 μm. (C’) FRAP graphs of Mid-Canoe stage wild-type (n=42) and *toga*-RNAi (n=20) spermatid. (C’’) Quantification of data in C’. Error bars are mean±s.d., ns = not significant, \*\*\**P*≤0.0003 (unpaired two-tailed t-test for A’, B’’, C’’).

These data provide a possible kinetic explanation for the mechanical failure observed in our fixed and live imaging. While Toga is not required for initial assembly of the perinuclear microtubule array, it does becomes important as these microtubules transition into a more stable scaffold during Mid-Canoe stages. We propose that as the mechanical strain on the sperm neck increases as the tissue develops, Toga acts to laterally cross-link (bundle) microtubules, or possibly cap the microtubule plus-ends, thereby reducing turnover. In the absence of this stabilization, microtubules remain more dynamic, leaving the sperm neck mechanically vulnerable and no longer able to reliably preserve a straight head-tail axis.

### Togaram is required for microtubule acetylation at the sperm head and neck

Given that microtubules in Mid-Canoe stage wild-type spermatids exhibited low dynamicity, we further investigated their stability by visualizing tubulin acetylation. Acetylation is a post-translational modification typically associated with long-lived, stable microtubules (Eshun-Wilson et al., 2019; Matsuyama et al., 2002; Piperno et al., 1987; Portran et al., 2017; Schulze et al., 1987; Xu et al., 2017). In *Drosophila* spermatids, the use of acetylated tubulin (ac-tubulin) as a marker for stable axonemal and cytoplasmic microtubules is well-established (Riparbelli et al., 2020). Based on the increased microtubule turnover observed in our FRAP experiments, we hypothesized that loss of Toga would result in a concomitant decrease in ac-tubulin levels.

In wild-type spermatids, we found that ac-tubulin surrounded the nucleus from Leaf stage (Figure 4A). Then, a dominant ac-tubulin bundle corresponding to the manchette formed on one side of the nucleus during Early Canoe stage (Figure 4A, orange arrows). Interestingly, we unexpectedly continued to observe a faint population of acetylated microtubules encircling the entire sperm head (Figure 4A, yellow arrow). As the spermatid progressed into Late Canoe and beyond, the manchette became less prominent, resulting in a more uniform acetylated microtubule distribution around the entire nucleus (Figure 4A, right panels). To distinguish this broader population of microtubules from the well-known manchette, we refer to them as the Sperm Microtubule Cage (SMC). Closer analysis reveals that in addition to encircling the nucleus, the SMC extends around and past the sperm neck region (Figure 4A’, Neck).

**Figure 4:**
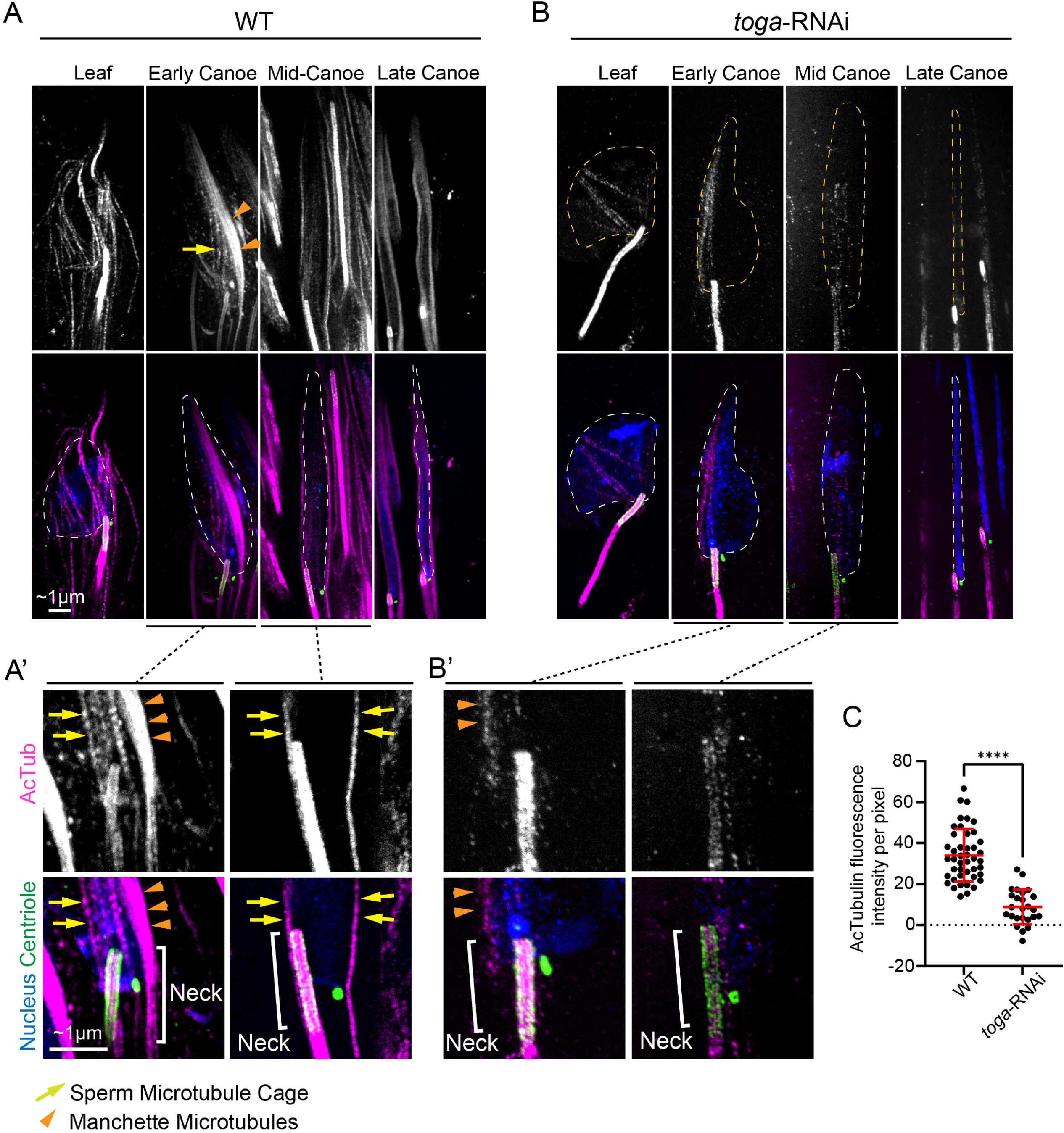
Togaram is required for proper levels of spermatid microtubule acetylation. (A,A’) Representative images showing wild-type spermatids during indicated developmental stages. Spermatids are labeled for the nucleus (DAPI, blue; dotted line), the centrioles (Ana1, green), and acetylated tubulin (magenta). Yellow arrow denotes Sperm Microtubule Cage (SMC). Orange arrowhead denotes manchette microtubules. Scale bar: ∼1 μm. (A’) Enlarged images of neck region from representative images. White bracket denotes neck region. (B,B’) Representative images showing *toga*-RNAi spermatids during indicated developmental stages. Spermatids are labeled for the nucleus (DAPI, blue; dotted line), the centrioles (Ana1, green), and acetylated tubulin (acTub, magenta). Scale bar: ∼1 μm. (B’) Enlarged images of neck region from representative images. White bracket denotes neck region. (C) Quantification of acetylated tubulin fluorescence intensity per pixel of wild-type (n=44) and *toga*-RNAi (n=25). Error bars are mean±s.d.; \*\*\*\**P*≤0.0001 (unpaired two-tailed t-test for C).

To test if Toga is required for microtubule acetylation, we performed comparative analysis on *toga-*RNAi spermatids. We found that in all stages, from Leaf to Late Needle, there was a drastic 75% reduction in ac-tubulin fluorescence at both the manchette and the SMC in the *toga*-RNAi sperm (Figure 4B,C). Furthermore, when specifically examining the neck microtubules, we observed a near 100% loss of ac-tubulin by the Mid-Canoe stage (Figure 4B’). This loss of stable microtubules in the neck region coincides with the developmental window when the off-axis phenotype manifests in *toga* loss-of-function spermatids. To assess the role of microtubule stability in spermatid alignment in an independent way, we treated wild-type testis with colchicine to depolymerize microtubules. We found that while 95% of control spermatids were linearly aligned, 55% of colchicine treated spermatids exhibited an off-axis phenotype (Figure S3A,B) with a highly variable attachment angle (Figure S3C). These data support a role for microtubules, and in particular stabile microtubules, in the alignment of the neck during spermiogenesis.

Taken together, our data reveal that Toga is required for robust acetylation of both manchette microtubules and the broader SMC population. These findings extend the microtubule scaffold relevant to sperm morphogenesis beyond the canonical manchette alone and suggest that the SMC contributes to neck mechanics as well. We therefore favor a model where Toga-mediated stabilization of the SMC and neck microtubules affords the structural rigidity necessary to withstand the mechanical forces and maintain proper sperm axial alignment during tissue development.

### Togaram binding partners Cep104 and CCDC66 are required for axial alignment

In mammalian cells, TOGARAM1 is a core component of the Ciliary Tip Module (CTM), a protein complex that localizes to the distal ends of cilia (Latour et al., 2020). The mammalian CTM comprises five proteins: CEP104, CSPP1, TOGARAM1, ARMC9, and CCDC66 (Latour et al., 2020). Biochemical and *in vitro* reconstitution studies have shown that the CTM functions as a specialized regulator of microtubule dynamics, effectively suppressing microtubule catastrophe and promoting slow, processive growth (Saunders et al., 2025). In *Drosophila*, three of the five CTM proteins are conserved (Figure 5A): Toga (CG42399), Cep104, and CG13251 (the putative ortholog of CCDC66), which we identified using the *Drosophila* Integrative Ortholog Prediction Tool (DIOPT) (Hu et al., 2011). Given Toga’s role in regulating spermatid microtubules, we hypothesized that Toga functions as part of a conserved *Drosophila* CTM.

**Figure 5:**
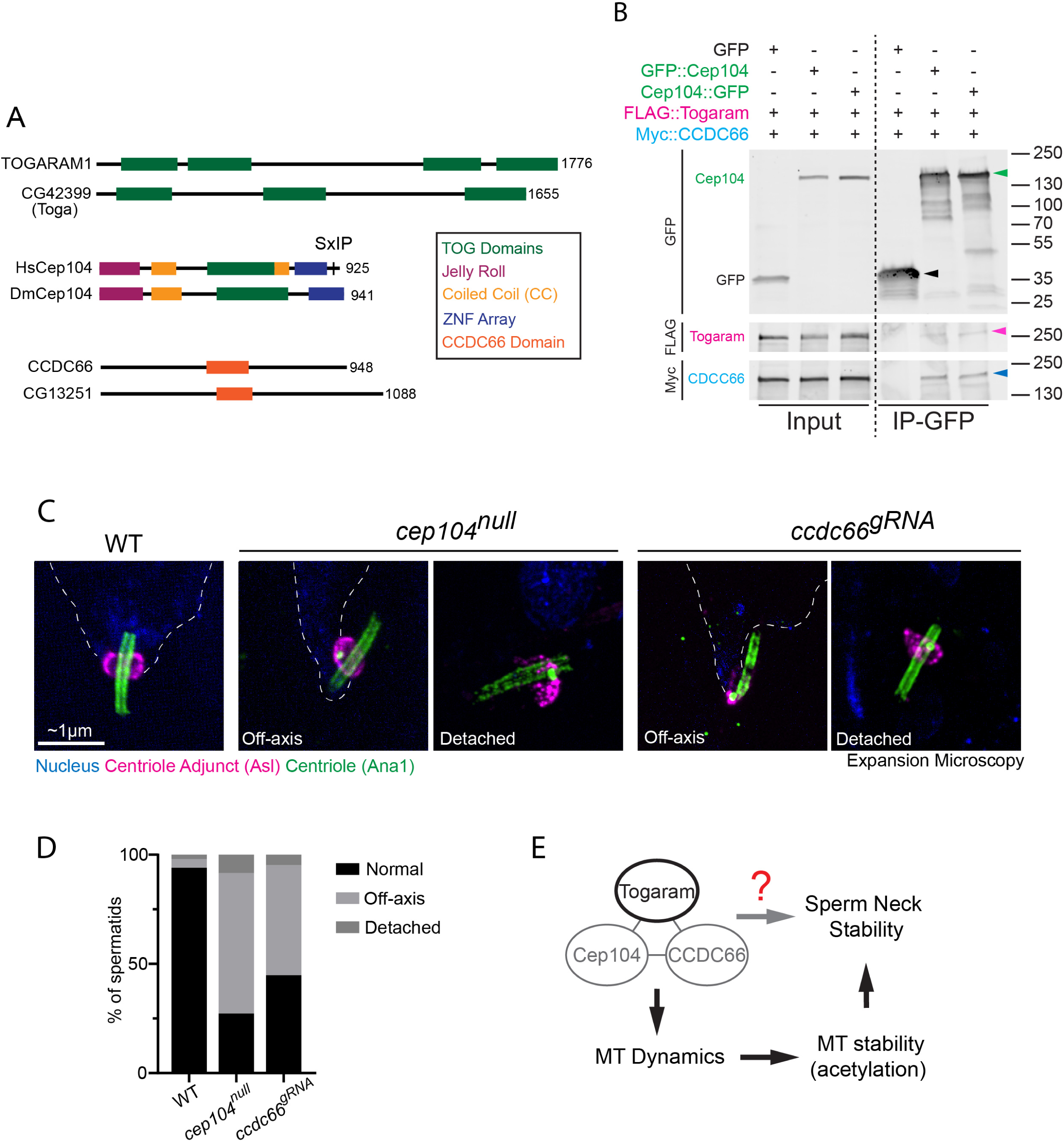
Togaram binding partners Cep104 and CCDC66 are required for sperm axial alignment. (A) Linear maps of human (top) and *Drosophila* (bottom) Togaram, Cep104, and CCDC66 depicting functional and structural domains. Domains are as indicated in the color key. (B) Both N– and C-terminally GFP-tagged Cep104 (green) binds Togaram (pink) and CCDC66 (blue). S2 cells were co-transfected with the indicated plasmids and anti-GFP co-IPs were prepared from cell lysates. Western blots of inputs and co-IPs were probed for GFP, Flag, and Myc. Arrow head colors correspond to indicated tagged protein colors. (C) Representative ExM images showing wild-type (left), *cep104^null^* (center), and *ccdc66^gRNA^* (right) Canoe stage spermatids with various alignment phenotypes. Spermatids were labeled for the nucleus (DAPI, blue; dotted line), the CA (Asl, magenta), and the centrioles (Ana1, green). Scale bar: ∼1 μm. (D) Quantification of wild-type (n=150), *cep104^null^* (n=98), and *ccdc66^gRNA^* (n=119) spermatids with various alignment phenotypes. (E) Working model in which Togaram, Cep104, and CCDC66 stabilize sperm head/neck microtubules to preserve a straight head-tail axis during the maintenance phase of spermiogenesis. Red question mark highlights a possible mechanism by which these proteins can directly regulate sperm neck stability independent of regulating microtubules.

We first tested whether these putative *Drosophila* CTM proteins physically interact. We co-overexpressed GFP::Cep104, Flag::Togaram, and Myc::CCDC66 in *Drosophila* S2 cells, and then showed that immunoprecipitation (IP) of GFP-Cep104 resulted in the robust co-IP of both Toga and CCDC66 (Figure 5B). This result indicates that Toga, Cep104, and CCDC66 can associate in *Drosophila* cells, consistent with a conserved complex analogous to the mammalian CTM. While these proteins might associate at microtubule plus-ends to regulate dynamics, it is equally plausible that they, individually or collectively, play roles independent of the CTM and microtubule plus-ends. For example, they could play regulatory roles by binding the microtubule lattice, or the microtubules of the centriole wall. Future studies will be essential to determine if these interactions are direct and whether they are specific to the sperm neck or can occur in other cell types.

As an initial test, however, we simply asked if Cep104 and CCDC66 might be required for axial alignment of the sperm neck. We performed loss-of-function experiments using a previously characterized null mutant of Cep104 (*cep104^null^*) (Ryniawec et al., 2023), or a germline specific gRNA targeting CCDC66 (*ccdc66^gRNA^*). We found that 64% of *cep104^null^* centrioles and 50% of *ccdc66^gRNA^* exhibited an off-axis phenotype, nearly identical to the loss of Toga (Figure 5C,D). Furthermore, 8% of *cep104^null^* and 5% of *ccdc66^gRNA^* centrioles were completely detached from nuclei (Fig. 5C,D).

These data identify Cep104 and CCDC66 as candidate components of the same axial-alignment pathway as Toga (Figure 5E). However, our data do not yet establish whether these proteins act as a single complex at the sperm neck, whether they regulate microtubule plus ends versus the lattice, or whether the mammalian CTM paradigm is fully conserved in spermatids. In our model, Toga, Cep104, and CCDC66 would stabilize sperm head and neck microtubules, promote their persistence long enough to accumulate stabilizing modifications such as acetylation, and thereby preserve a straight head-tail axis during the maintenance phase of spermiogenesis (Figure 5E).

Our findings support a model in which Togaram acts during the maintenance phase of spermiogenesis to stabilize sperm head and neck microtubules, thereby preserving a straight head-tail axis as the spermatid undergoes extensive remodeling. In doing so, this work separates the initial establishment of head-tail attachment from the later mechanical reinforcement needed to prevent neck buckling and maintain axial alignment. The identification of the Sperm Microtubule Cage and the shared requirement for Cep104 and CCDC66 further suggest that a conserved microtubule-regulatory module underlies this process. Future studies should define how these factors act at the sperm neck, whether through direct regulation of microtubule plus ends, lattice stabilization, or higher-order bundling, and test the extent to which this mechanism is conserved in mammalian spermiogenesis and male infertility.

## METHODS AND MATERIALS

### Flies

Fly stocks were maintained on Bloomington-recipe fly food (LabExpress). Crosses were performed at 25°C. The following fly strains were used in this study: y,w (gift from Mark Peifer, University of North Carolina, Chapel Hill), CG42399::GFP (Bloomington Drosophila Stock Center, Bloomington, IN, USA, #65336), PACT::GFP (gift from Jordan Raff, University of Cambridge, Cambridge), CG42399 RNAi (Vienna Drosophila Resource Center, Vienna, Austria #110484), bam-Gal4 (Bloomington Drosophila Stock Center, #80579), ubi-GFP::tubulin (gift from Tomer Avidor-Reiss, University of Toledo) H2A::RFP (Bloomington Drosophila Stock Center, #23651), cep104^null^ (Ryniawec et al., 2023), CG13251 sgRNA (Bloomington Drosophila Stock Center, #85911), vas-Cas9 (Bloomington Drosophila Stock Center, #51324).

### Testis fixation and immunofluorescence

Testes were dissected in Schneider’s medium with antibiotic-antimycotic (S2 media) and fixed in 9% paraformaldehyde (PFA) at room temperature (RT) for 15-20 minutes. Testes were washed in PBS + 0.3% Triton X-100 (PBST) then blocked for 1-2 hours at RT in PBST with 5% normal goat serum (NGS) (MilliporeSigma, Burlington, MA, USA). Samples were incubated in primary antibody in blocking solution at 4°C overnight (guinea pig anti-Asterless 1:10,000, mouse monoclonal anti-polyglutamylated tubulin GT335 1:500, rat anti-Togaram 1:50). Samples were then washed 3x 10 minutes in PBST and incubated in secondary antibody in blocking solution for 4 hours at RT (1:1000). DAPI staining was performed concurrently with the secondary antibody (1:50,000). After washing 3x 10 minutes in PBST, samples were mounted in Aqua-Poly/Mount (Polysciences) and covered with a #1.5 coverslip for confocal imaging.

For analysis of tubulin fluorescence intensity, sample preparation deviated slightly and included squashing of tissue. After dissection, an individual testis was moved to a drop of S2 media on a clean slide. Using forceps, the testis was opened at the midway point. A #1.5 coverslip was placed on the media drop, and media was removed using a Kimwipe (Kimberly-Clark, Irving, TX, USA) until tissue appeared squashed. Slides were flash frozen in liquid nitrogen, coverslip removed using a razor blade, and incubated in 100% methanol at –20°C for at least 20 minutes. Samples were then blocked for 30 minutes in 5% NGS in PBS + 0.1% Tween20 (PBST). Samples were incubated in primary antibody in blocking solution at RT for 1 hour followed by 3x 10 minute washes with PBST. Samples were then incubated in secondary antibody with DAPI in blocking solution at RT for 1 hour and slides were then dunked in 1XPBS 10 times. Samples were mounted as stated above.

### Confocal imaging

Confocal images were acquired using a Nikon Eclipse Ti2 (Nikon Instruments, Melville, NY, USA) with a Yokogawa CSU-W1 spinning disk confocal head (Yokogawa Life Science, Sugar Land, TX, USA) equipped with a Prime BSI CMOS camera (Teledyne Photometrics, Tucson, AZ, USA) and Nikon Elements Software (Nikon Instruments). Non-expanded samples were imaged using a 100xTIRF/1.49 NA oil immersion objective, while expanded samples were imaged using a 100x/1.35 NA silicone immersion objective. For analysis of mislocalized DNA aggregates, a 10x/0.3 NA air objective was used. For analysis of seminal vesicle area, a 20x/0.75 NA air objective was used. 405-, 488-, 568-, and 647-nm laser lines were used for all image acquisition. All images were analyzed and processed in FIJI (ImageJ, National Institutes of Health, Bethesda, MD, USA).

### Expansion microscopy

Expansion microscopy (ExM) on *Drosophila* testis tissue was carried out as described in (Burns et al., 2026). Testes were dissected and fixed in anchoring solution (FA/AA) at 37°C overnight. Humid chamber was prepared and testes were incubated in Monomer solution (MS; 500 µl 38% sodium acrylate solution, 250 µl 40% acrylamide, 100 µl 10× PBS, 50 µl 2% *N*,*N′*-methylenebisacrylamide) below a coverslip for 1 hour at 37°C for gel polymerization. Gels were biopsy punched and incubated at RT for 15 minutes and 95°C for 1.5 hours in denaturation buffer (10 ml 0.5 M Tris pH 9.0, 4 ml 5 M NaCl and 57.14 ml 10% SDS). At RT, gels were expanded in H_2_O followed by two washes in 1X PBS before primary antibody staining at RT for 24 hours (chicken anti-Ana1 1:1000, guinea pig anti-Asterless 1:5000, mouse monoclonal anti-acetylated tubulin 1:100, rat anti-Togaram 1:50). Gels were washed 3x 10 minutes with PBS 0.1% Tween20 before secondary antibody staining at RT overnight (secondaries 1:500, DAPI stain 1:25,000). Gels were fully expanded in H_2_O at RT and mounted on a poly-L-lysine-coated glass-bottom dish for imaging as stated above.

### Live-tissue imaging

Testes were dissected and mounted in a drop of S2 media, surrounded by Halocarbon oil 700 (Sigma-Aldrich) on a 50-mm lummox dish (Sarstedt), and covered with a #1.5 coverslip. Live images were acquired on the same microscope as fixed images with a 100xTIRF/1.49 NA oil immersion objective. Images were collected at 1 minute intervals.

### Fluorescence recovery after photobleaching (FRAP)

Testes were dissected and mounted in S2 media and covered with a #1.5 coverslip. FRAP was performed on the same microscope as fixed images with a 100xTIRF/1.49 NA oil immersion objective. Bleaching was performed with a rectangular region of interest (ROI) using the 405-nm laser operating at 5% laser power. A single iteration was used for the bleach pulse which lasted 10ms. Fluorescence recovery was monitored with 488-nm laser line at 25% laser intensity and 150ms exposure. Images were captured at 1s intervals and tissue was recorded a minimum of 10s before bleaching and 60s after.

### Qualitative Localization Screen

Screen list was assembled from genes within the fly sperm proteome with publicly available GFP lines (Bloomington Drosophila Stock Center). From this list, priority was given to microtubule binding proteins and unknown proteins. For each line, testes were immunostained and imaged as stated above. A positive localization result was defined as visualization of the protein at/near the sperm neck.

### Seminal vesicle area analysis

Whole male reproductive tracts were dissected in S2 media and fixed in 9% paraformaldehyde (PFA) for 15-20 minutes. Samples were stained with DAPI 405 for 3 hours at RT. Seminal vesicles were further dissected from the rest of the reproductive tract, mounted in Aqua-Poly/Mount (Polysciences), and covered with a #1.5 coverslip for confocal imaging. Seminal vesicles were imaged using a 20x/0.75 NA air objective and area was quantified as determined by DAPI staining.

### Fertility assay

Fertility assays were conducted on four distinct genotypes of male flies. Three groups were control genotypes: y,w, BamGal4, and *toga*-RNAi, and one group was the experimental genotype: BamGal4>*toga*-RNAi. For each group 10-20 vials were set up containing two y,w virgin females with one naïve male of the respective genotype. Virgin females and naïve males were allowed to mate for 3 days at 25°C. After the mating period, the parental generation was removed. All adult progeny were counted for 10 days following the first eclosion.

### Analysis of sperm neck alignment

Leaf and Early Canoe stage spermatids were selected for analysis. Nonexpanded samples were measured. Spermatids were only analyzed if they appeared parallel to the imaging plane and the nuclei (DAPI) and centriole (PACT::GFP) could be clearly identified. Spermatids were scored as ‘Normal’ if the sperm neck angle was 160-180°. Angles less than this range were scored as ‘Off-axis’. Spermatids were scored as ‘Detached’ if the centriole was not in contact with the nucleus. From this analysis, the sperm neck angle and percent of normal, bent, and detached centrioles is presented.

### Analysis of properly localized spermatids per cyst

To determine if the absence of off axis spermatids at later stages of spermiogenesis was a result of axis correction or spermatid elimination, we analyzed cysts of Mid– and Late Canoe stage spermatids to determine how many spermatids remained in the proper location. A rectangular ROI of approximately ∼1000µm^2^ was drawn around the basal end of each cyst to include the nuclei clustered there. Using PACT::GFP-labeled centrioles as a proxy for spermatid number, the number of spermatids in this ROI was counted. Since it is possible that centrioles may be detached from nuclei in the experimental genotype, this analysis would not reveal decapitated spermatids whose nuclei and centrioles were not displaced from the most basal section of the cyst. The reported numbers therefore likely underestimate the extent of spermatid disorganization and elimination within an indicated cyst. The number of basally localized centrioles per cyst is presented.

### Analysis of tubulin intensity

Testes expressing GFP::tubulin were lightly squashed under a coverslip and imaged live. Only cysts of spermatids from the Early to Mid-Canoe stage were analyzed. Multiple rectangular regions of interest (ROIs) were drawn at the spermatid heads of a cyst. The mean fluorescence of GFP::tubulin at the spermatid heads within each cyst was calculated. Background subtraction was applied by measuring average background intensity obtained from multiple ROIs drawn in regions surrounding the cyst, then subtracting this from the mean cyst intensity.

### Analysis of FRAP

Only spermatids that remained in frame for the duration of the experiment were analyzed. Spermatids were binned by Early Canoe and Mid-Canoe and analysis was conducted at both stages. To measure recovery, an ROI was drawn within the bleached area of the spermatid head and the raw integrated density of ubi-GFP::tubulin fluorescence was measured every frame from 10 frames prior to bleaching to 30 frames after bleaching. An measurements of an ROI of the same size in the adjacent background were taken and subtracted. To correct for photobleaching from the imaging laser, the signal at each timepoint was divided by fluorescent measurements from an unaffected region of the spermatid heads at that time timepoint. With this data corrected for background fluorescence and normal photobleaching, the pre-bleach intensity was then set to 1.0 and bleached intensity as 0. This provides the relative fluorescence recovery curve after bleaching for each spermatid analyzed. The mobile fraction was determined by fitting this curve using Prism (Y=Y0 + (Plateau-Y0)*(1-exp(-K*x)) –“one-phase association” – GraphPad, Boston, MA, USA)

### Analysis of acetylated tubulin intensity

Expansion microscopy images of Canoe stage spermatids with nuclei parallel to the imaging plane were analyzed. An ROI of 15×50 pixels was drawn around the sperm head. The acetylated tubulin average fluorescence intensity per pixel was measured. An ROI of the same size was drawn around the adjacent background, the fluorescence intensity per pixel was measured, and then subtracted from the sperm head measurement. The acetylated tubulin fluorescence intensity per pixel is presented from this analysis.

### Cell culture

*Drosophila* S2 cells (DGRC Stock 181; https://dgrc.bio.indiana.edu//stock/181; RRID:CVCL_Z992) were maintained in Schneider’s *Drosophila* medium (ThermoFisher Scientific, Waltham, MA) supplemented with 10% heat-inactivated fetal bovine serum (ThermoFisher Scientific or Cell Culture Collective) and 1× Antibiotic-Antimycotic (ThermoFisher Scientific) at 27°C. Cells were transfected using Effectene (Qiagen, Germantown, MD); transfections were performed using 1µg of each plasmid and 4 × 10^6^ cells as directed by the manufacturer.

### DNA constructs

A multiple cloning site (MCS) was generated by PCR amplification (Phusion polymerase, Thermo Fisher Scientific) of overlapping oligos and TOPO cloned into pENTR to generate pENTR-MCS. Genomic DNA of CG13251 was PCR amplified from y,w flies and TOPO cloned into pENTR vector to make pENTR-CG13251. Full-length cDNA of Cep104 (Rogers lab, University of Arizona) was PCR amplified and TOPO cloned into pENTR vector to generate pENTR-Cep104. Togaram was synthesized as two pieces from Twist Biosciences (South San Francisco, CA. The two pieces were then PCR amplified and cloned using Gibson assembly into pENTR-MCS vector to generate pENTR-Togaram. S2 cell expression vectors were generated by the Gateway cloning system (ThermoFisher Scientific) to create vectors with Actin5C promoter and the tags of interest (pAFHW, pAGW, pAMW, pAWG; F = Flag, G = GFP, M = Myc; https://emb.carnegiescience.edu/drosophila-gateway-vector-collection). pAGW-MCS plasmid, expressing GFP alone was used as GFP control for IP experiments.

### Antibody production and validation

Bacterial construct cloning, bacterial protein expression, protein purification, and antibody generation from three Wistar rats against amino acids 1-664 of Togaram was performed by GenScript USA, Inc. (Piscataway, NJ). All antibodies were tested in *Drosophila* testes by immunostaining, Rat #1 showed the expected staining patterns and was used for all subsequent experiments.

### Immunoprecipitations

Rabbit anti-GFP antibody (Abcam Inc., Cambridge, MA) was conjugated to Protein-A Dynabeads (Thermo Fisher Scientific) for 1 hour at 4°C with mixing. Cells were harvested 2 days after transfection and resuspended in lysis buffer [50 mM Tris pH 7.4, 150 mM NaCl, 0.5% Triton X-100, 1 mM Dithiothreitol (Fisher Scientific, Pittsburgh, PA), cOmplete™, EDTA-free Protease Inhibitor Cocktail (Sigma-Aldrich, St. Louis, MO)]. After 5 minutes on ice, the lysate was cleared for 5 minutes, 17,200× g, 4°C. The supernatant was then incubated with the conjugated GFP-Protein A Dynabeads for 1 hour at 4°C with mixing. The GFP tagged protein bound to the Dynabeads was washed thrice in lysis buffer on ice, each wash for 5 minutes, eluted by boiling in 2× SDS-sample buffer [58-mM Tris pH 6.8, 5% glycerol, 1.95% SDS, 1.55% DTT, 0.05% Bromophenol Blue] for 5 minutes, and stored at −80°C until use in Western blotting. The input lysate was also analyzed.

### Western blots

Samples were run on 4-20% SDS–polyacrylamide gradient gels. Samples were transferred to Polyvinylidene fluoride [PVDF] membrane using Tris-Glycine transfer buffer (Thermo Fisher Scientific) with 20% methanol. Blots were blocked in 5% nonfat dry milk diluted in TBST [0.1% Tween 20 diluted in Tris Buffered Saline—50-mM Tris-HCl, pH7.5, 150-mM NaCl] for 60 minutes before incubation with primary antibodies diluted in blocking solution overnight at 4°C. Primary antibodies were anti-Flag (M2, 1:1000, Sigma Aldrich), anti-GFP (JL8; 1: 5000; Takara Bio USA, Mountain View, CA), anti-myc (E4F4D, 1:1000, Cell Signaling Technology, Danvers, MA); Blots were washed in TBST, the blots were incubated in secondary antibodies diluted in blocking solution for 1 hour at RT. Secondary antibodies were Alexa Fluor 647 goat anti-mouse IgG1, Alexa Fluor 488 goat anti-mouse IgG2a, Alexa Fluor 568 goat anti-mouse IgG2b (all 1:5000; ThermoFisher Scientific). The blots were then washed and detected using an Amersham™ ImageQuant™ 800 Western blot Imaging System (Cytiva, Marlborough, MA).

### Statistics

The following statistical test was performed: two-tailed, unpaired t-test (Figures 2B’, 2C’, 2D’, 3A’, 3B’’, 3C’’, and 4C) and one-way ANOVA with Tukey’s correction (Figure 2D’’).

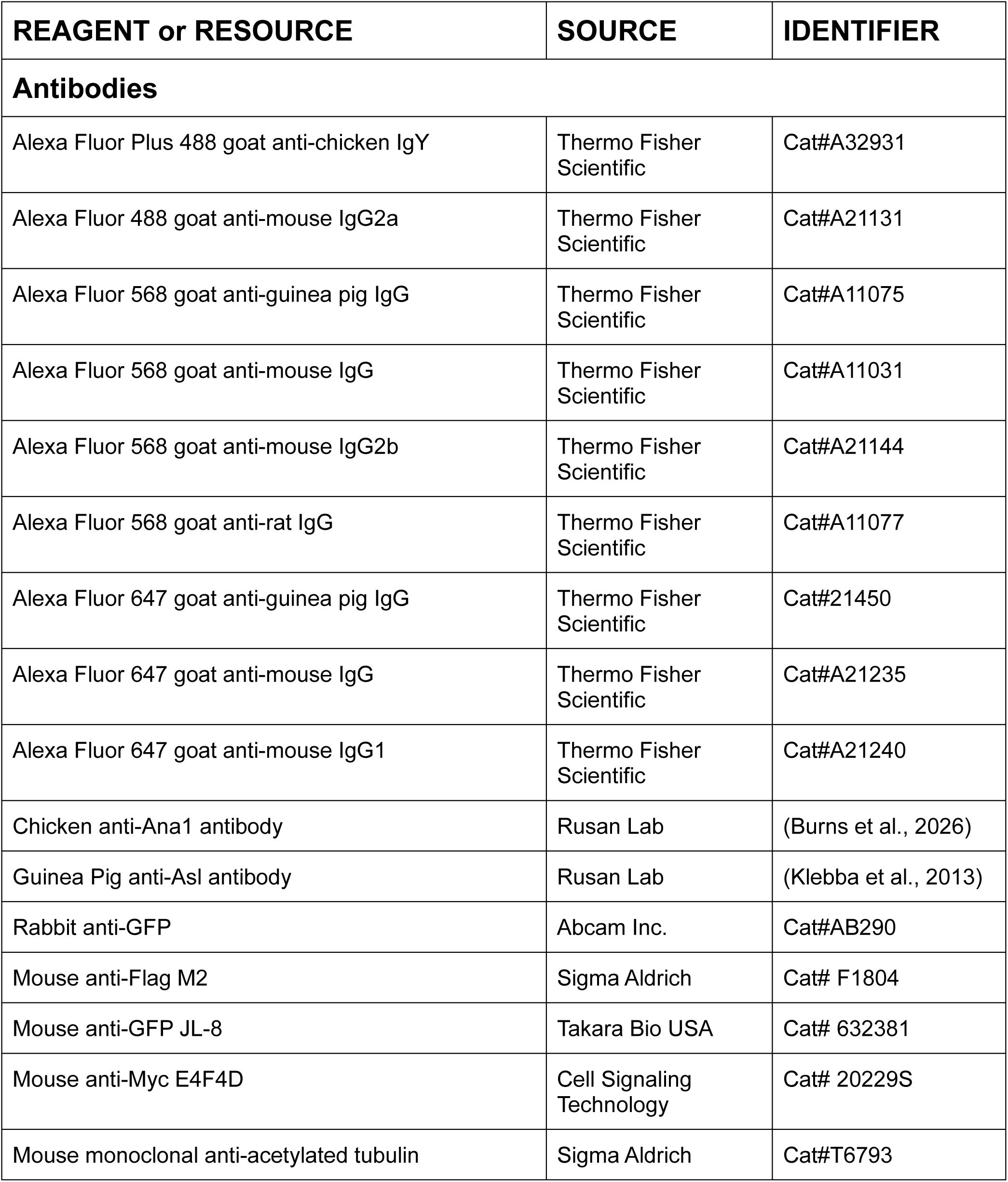
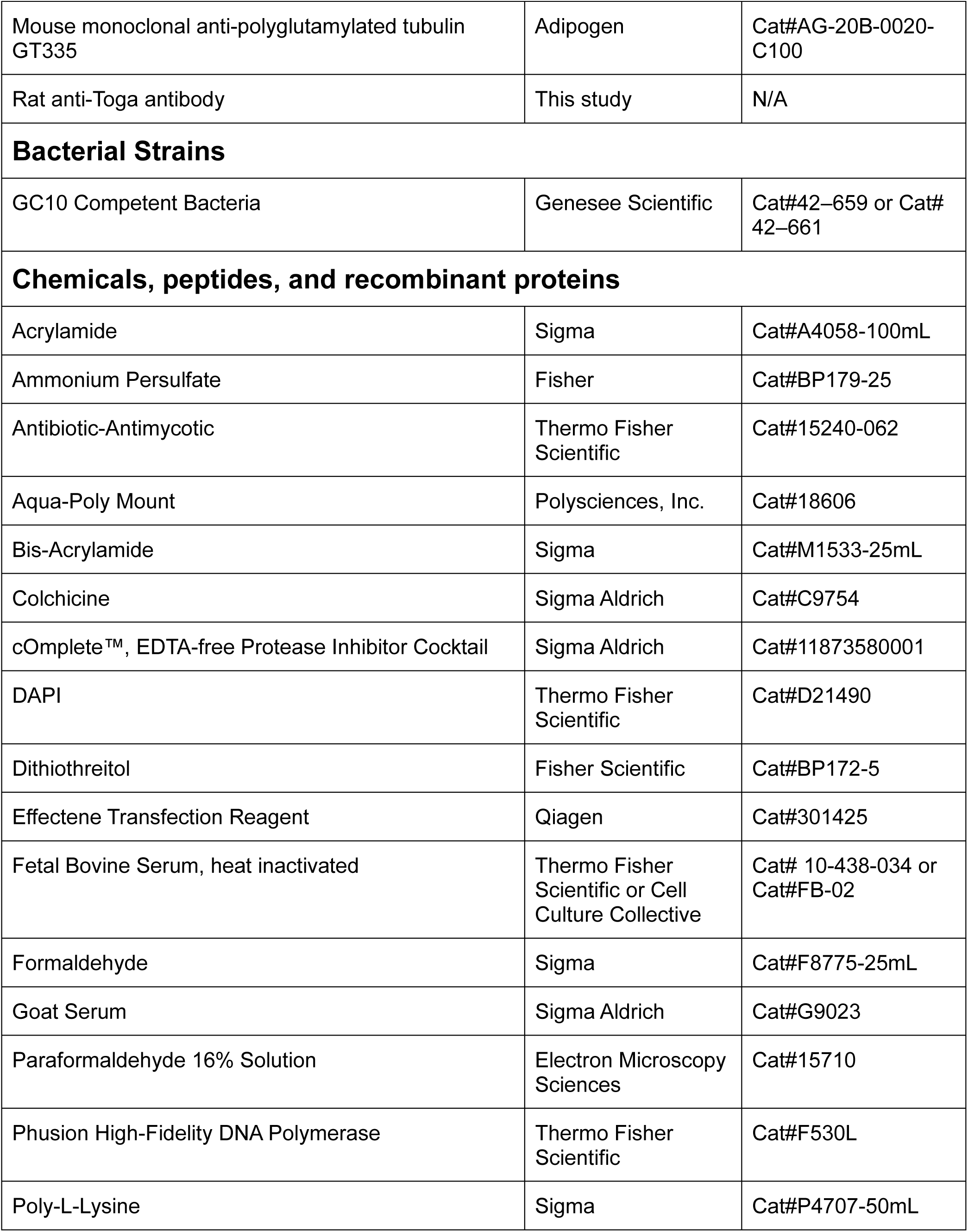
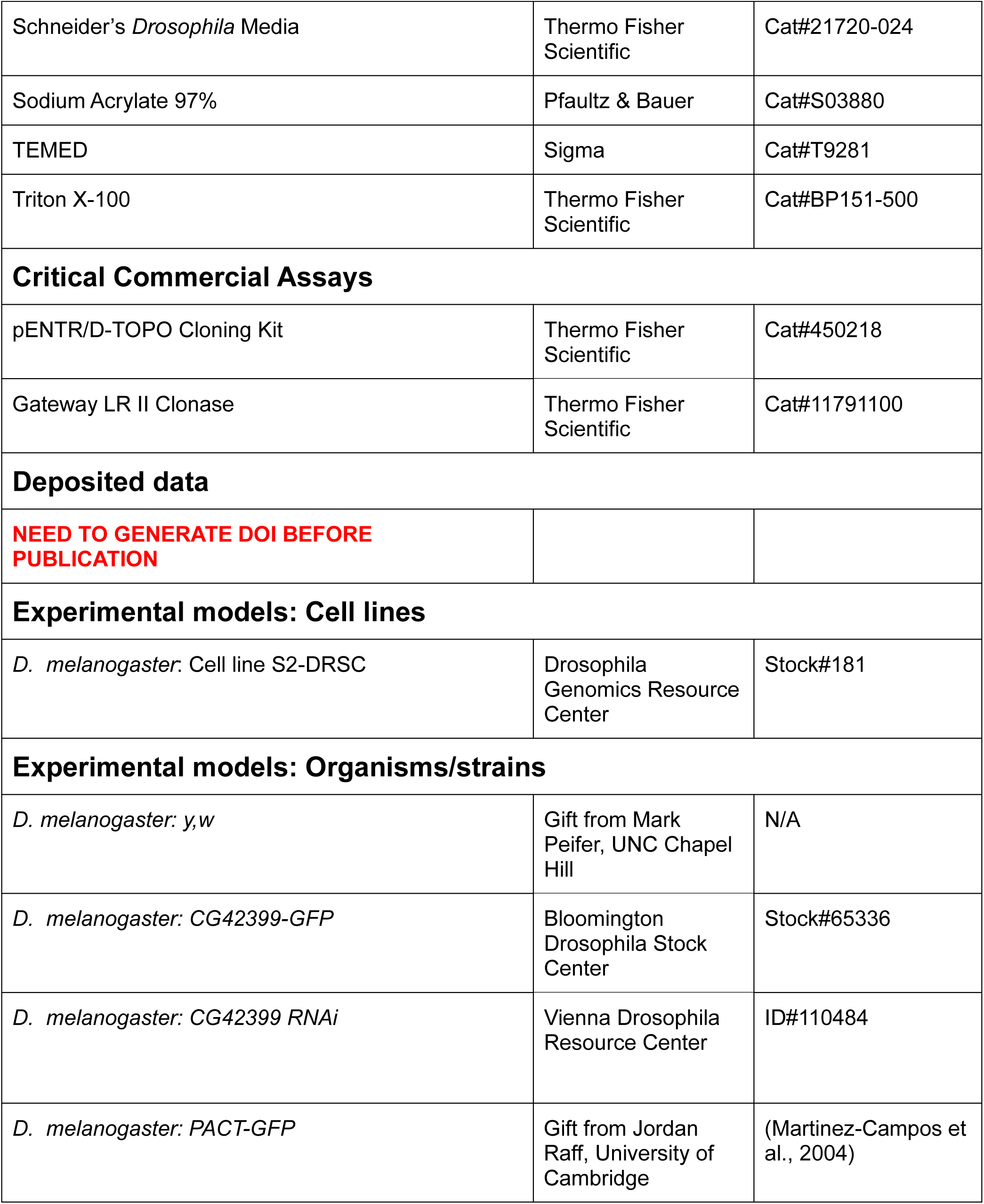
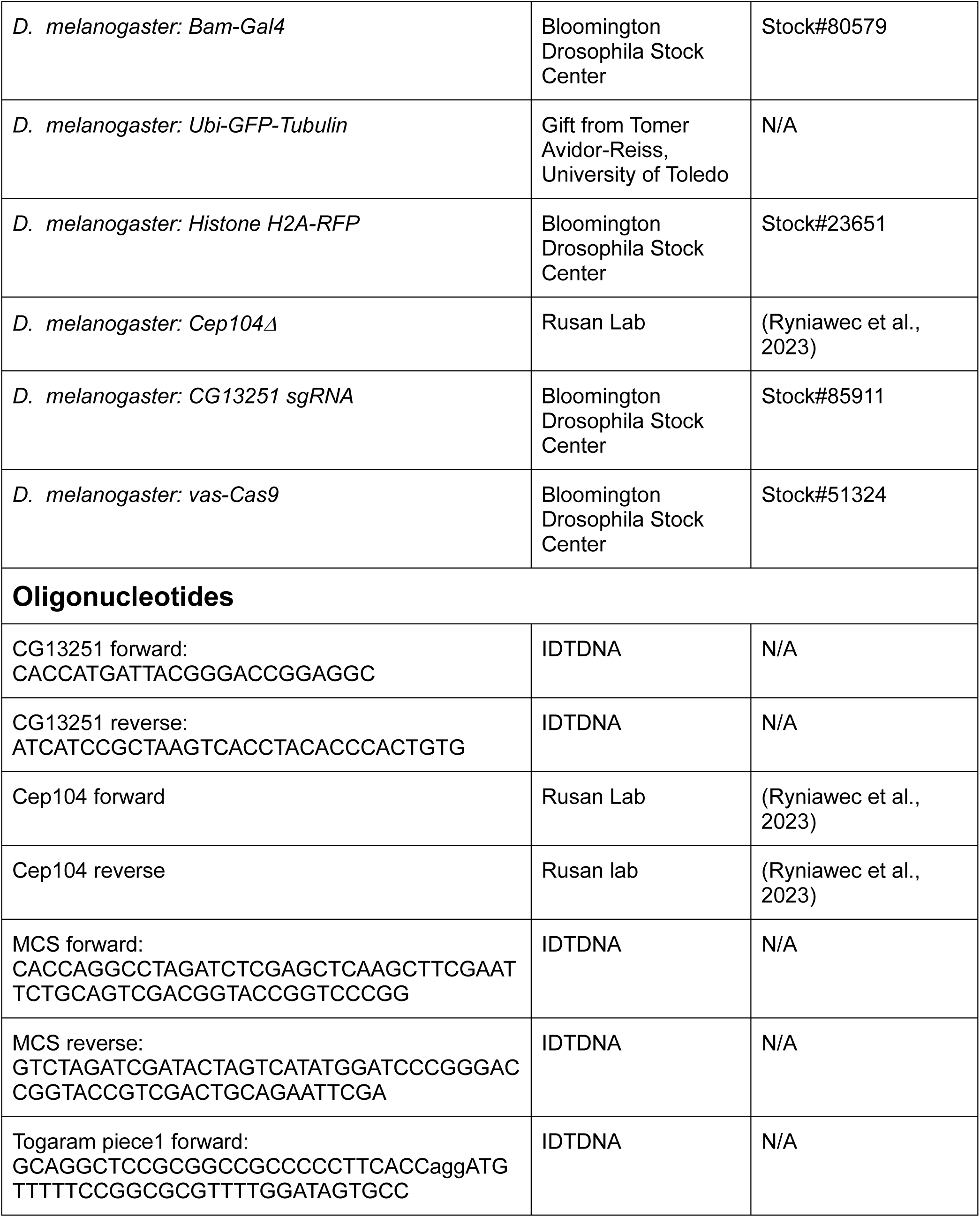
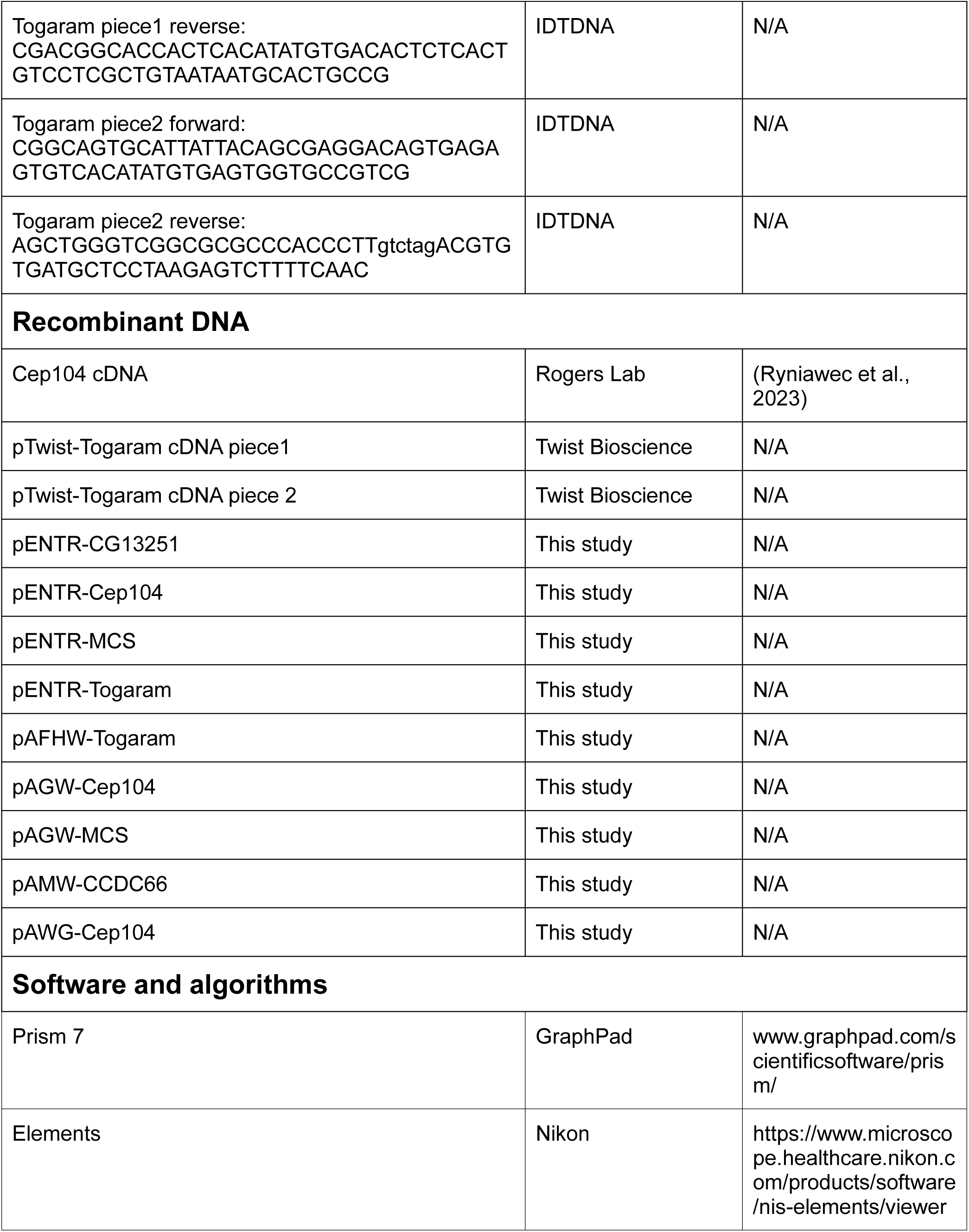
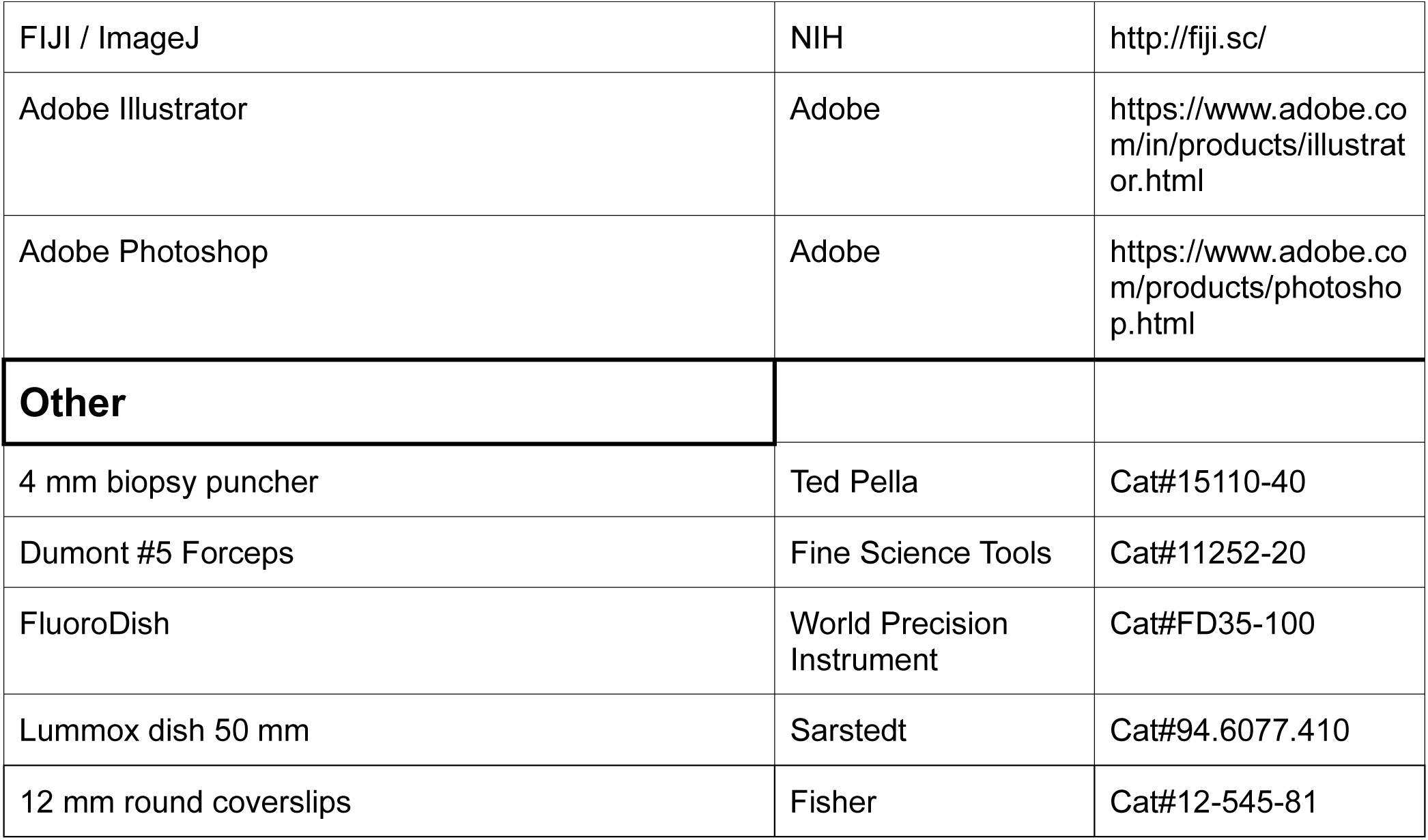
TABLE 1.

## Acknowledgements

We thank Brian Galletta for help with microscopy and data quantification and Danielle Buglak for her mentorship and guidance. We also thank Rusan lab members for helpful discussion. This work was supported by the Division of Intramural Research at the National Heart, Lung, and Blood Institute (ZIAHL006126 to NMR).

## Disclaimer

The contributions of the NIH author(s) are considered Works of the United States Government. The findings and conclusions presented in this paper are those of the author(s) and do not necessarily reflect the views of the NIH or the U.S. Department of Health and Human Services.

## Data availability statement

All relevant data can be found within the article and its supplementary information. The DOI for all raw data is XXXXXXXX

---

**Figure S1:**
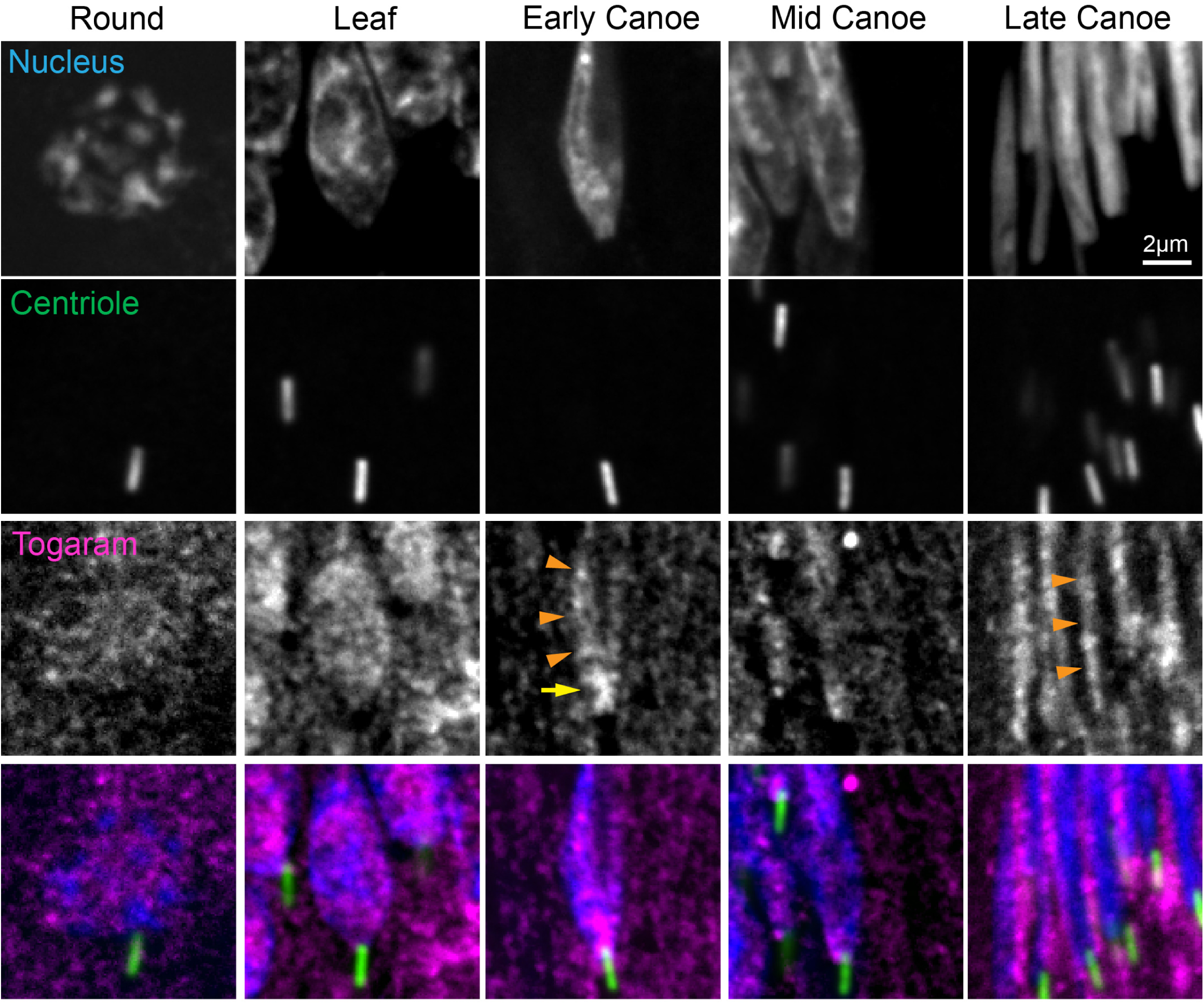
Togaram dynamically localizes to sperm nucleus and neck. Representative images showing wild-type spermatids during indicated developmental stages. Spermatids are labeled for the nucleus (DAPI, blue), centriole (PACT::GFP, green), and Togaram (Toga, magenta). Yellow arrow denotes neck localization. Orange arrowheads denote nuclear localization. Scale bar: 2 μm.

**Figure S2:**
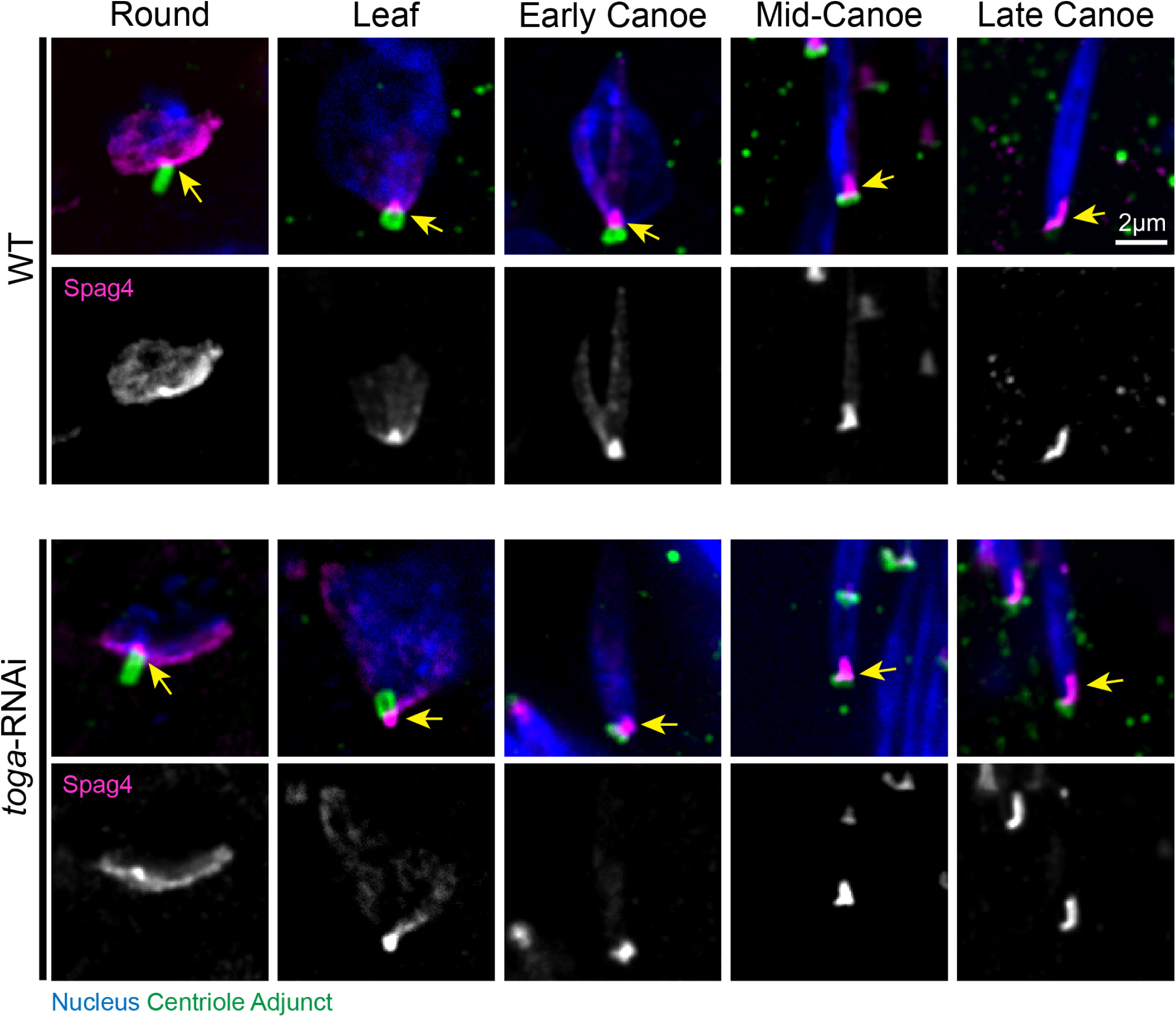
Togaram does not disrupt Spag 4. Representative images showing wild-type (top) and toga-RNAi (bottom) spermatids during indicated developmental stages. Spermatids are labeled for the nucleus (DAPI, blue), the centriole adjunct (Asl, green), and Spag4 (magenta). Yellow arrows indicate the HTCA. Scale bar: 2 μm.

**Figure S3:**
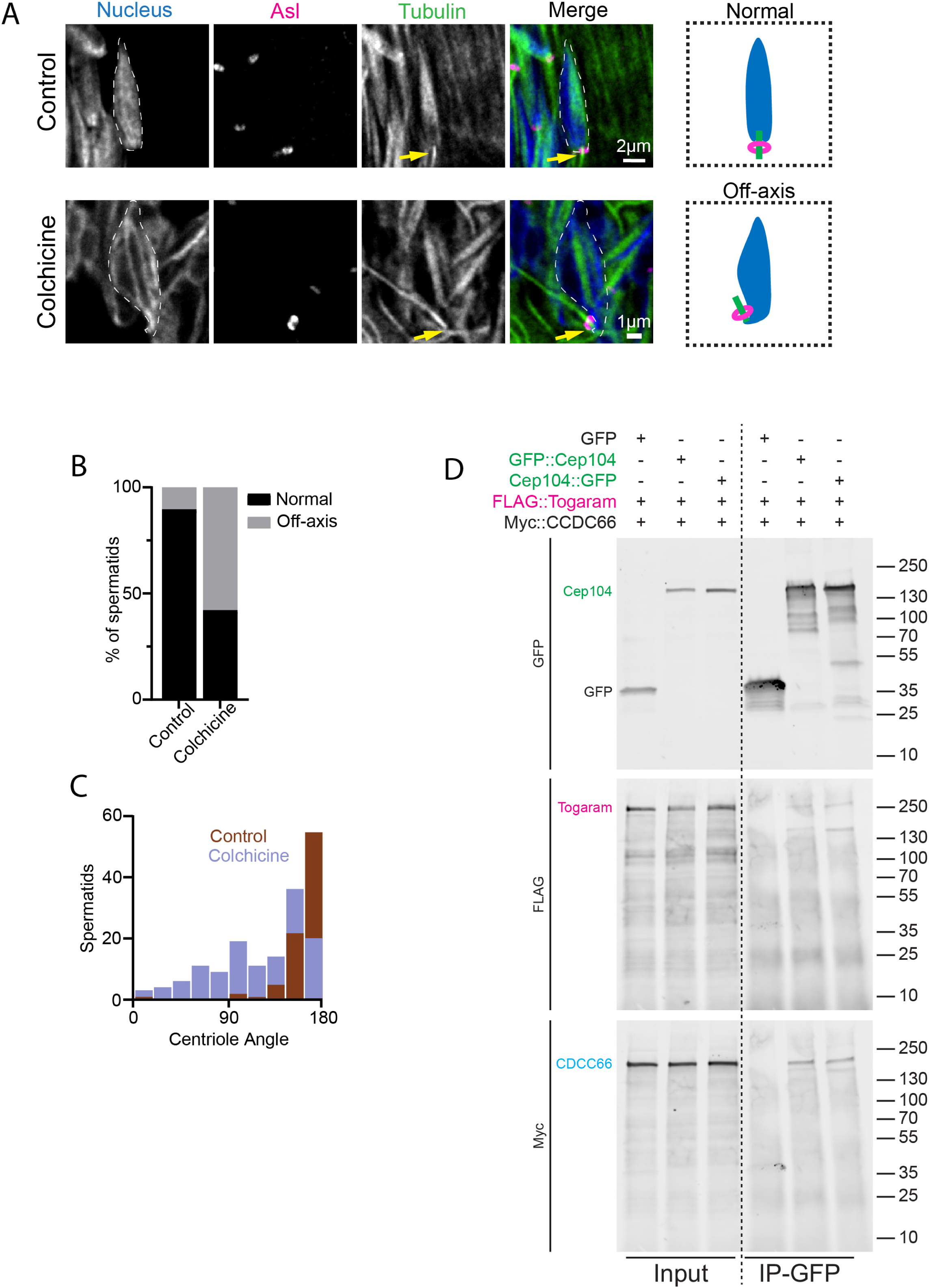
Stable spermatid microtubules are required for sperm neck alignment. (A) Representative images showing control (top) or colchicine treated (bottom) Canoe stage spermatids. Spermatids are labeled for the nucleus (DAPI, blue), CA (Asl, magenta), and tubulin (ubi-GFP::tubulin, green). Yellow arrow indicates neck region. Cartoons depict spermatids with centrioles that are Normal or Off-axis in relation to nucleus. Scale bar: 2 μm (top), 1μm (bottom). (B) Quantification of control (n=86) and colchicine treated (n=133) spermatids with various alignment phenotypes. (C) Quantification of attached centriole angle to nucleus in control (n=86) and colchicine treated (n=133). (D) Raw gel images corresponding to Figure 5B. Both N– and C-terminally GFP-tagged Cep104 (green) binds Togaram (pink) and CCDC66 (blue). S2 cells were co-transfected with the indicated plasmids and anti-GFP co-IPs were prepared from cell lysates. Western blots of inputs and co-IPs were probed for GFP, Flag, and Myc.

**Table S1.** Qualitative GFP localization screen.

| **Stock #** | **Gene Name** | **Localization** |
| --- | --- | --- |
| 59974 | CG10462 | No significant signal |
| 59975 | CG10631 | No significant signal |
| 68659 | CG10979 | No significant signal |
| 56785 | CG12155 | No significant signal |
| 66590 | CG1233 | Head of mature sperm |
| 92384 | CG14441 | Head of mature sperm |
| 60235 | CG1632 | Head of mature sperm |
| 6843 | CG1640 | Mitochondria |
| 50877 | CG1677 | Head of mature sperm |
| 63153 | CG17349 | Tails of mature sperm |
| 67660 | CG2199 | Nuclei of spermatogonia |
| 81263 | CG2712 | Head of mature sperm |
| 60175 | CG31183 | No significant signal |
| 60148 | CG3339 | Distal end of round stage spermatids centrioles |
| 61775 | CG34357 | Head of mature sperm |
| 50826 | CG3939 | No significant signal |
| 65336 | CG42399 | Sperm neck region |
| 92370 | CG4496 | No significant signal |
| 50855 | CG8036 | No significant signal |
| 50834 | CG8209 | Cytoplasm of spermatocytes |
| 50802 | Cindr | Muscle tissue and individualization cones |
| 60190 | Clip-190 | Nuclear envelope of round stage spermatids and tails |
| 58447 | Don juan | Tails of sperm |
| 60193 | Karst | No significant signal |
| 58406 | Protamine B | Head of elongating and mature sperm |
| 39647 | Rhea | Head of mature sperm |
| 91747 | Spc105R | No significant signal |
| 59827 | Spire | No significant signal |
| 60258 | Sprint | No significant signal |
| 60525 | Obscurin | No significant signal |

